# Relationship between SARC-F score and Peripheral Blood Lymphocytes in Juvenile Idiopathic Arthritis: An Observational Pilot Study

**DOI:** 10.1101/2025.10.16.681879

**Authors:** Zahra Ladan, Amaani Ahmad, Sameera Oruganti, Souraya Sayegh, Charalambos Hadjicharalambous, Nicole Zhang, Madhura Castelino, Corinne Fisher, Maria Leandro, James Glanville, Arne Akbar, Venkat R Reddy, Debajit Sen

## Abstract

**Background:** Given the relatively high prevalence of sarcopenia and immune profile abnormalities in inflammatory arthritis such as rheumatoid arthritis (RA), we investigated the relationship between exercise habits, SARC-F score and peripheral blood lymphocyte profiles in patients with Juvenile idiopathic arthritis (JIA).

**Methods:** All participants with JIA completed the SARC-F survey and self-reported exercise habits through online questionnaires. We defined two groups based on SARC-F scores as musculoskeletal function impaired (SARC-F^hi^+) and musculoskeletal function not impaired (SARC-F^hi^-). We used flow cytometry of peripheral blood mononuclear cells to define immune cell subpopulations.

Statistical analyses were performed using GraphPad Prism software version10.

**Results:** Of the 18 participants with a median age of 24 years – five belonged to the SARC-F^hi^+ group and 13 to the SARC-F^hi^-group, of whom 80% and 38% were female, respectively. The SARC-F^hi^ + group had fewer CD45RA+CD27+ naïve CD4+ T cells (26.60% vs. 45.30%, *p=0*.*0264*) and greater CD45RA-CD27-effector memory CD4+ T cells (11.90% vs. 7.00%, *p=0*.*0350*). The SARC-F^hi^+ group also had fewer CD56-CD16+ NK cells compared to the SARC-F^hi^-group (2.60% v. 4.28%, *p = 0*.*0194*). The moderate exercise group (n = 13) had increased numbers of total CD4+ T cells (54.06% vs. 37.30%, *p=0*.*0140*) and 3-fold fewer CD45RA+CD27-EMRA CD8+ T cells (3.17% vs. 10.40%, *p=0*.*0350*). There were no other significant differences for other immune cells / subpopulations including B cells.

**Conclusions:** Our study findings suggest notable differences in immune profile relative to SARC-F score. Furthermore, the 3-fold lower frequency of CD8+ TEMRA subpopulation of T cells, reflective of immunosenescence, suggest a potential inverse relationship between moderate exercise and peripheral immune cell profile in JIA.

**Key messages:**

1. SARC-F score was used as a clinically relevant tool to identify musculoskeletal function in JIA
2. SARC-F^hi^+ group had fewer naïve CD4+ T cells and greater effector memory CD4+ T cells
3. Moderate exercise group had 3-fold lower frequency of CD8+ TEMRA T cells

**Graphical Abstract:** 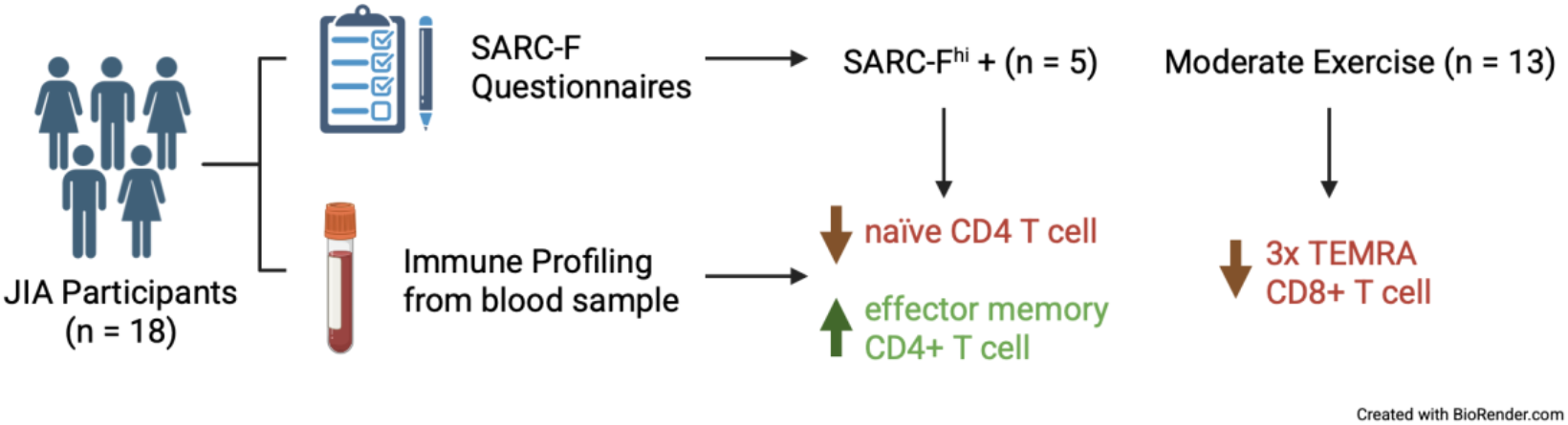

## Introduction

A complex, bidirectional relationship between the skeletal muscle and the immune system influences chronic inflammation in ageing (1) and autoimmunity (2). The progressive loss of skeletal muscle mass, strength and function reflecting a generalised disorder of skeletal muscle is referred to as sarcopenia (3, 4) . Sarcopenia is associated with poorer quality of life and increased risk of falls, fractures and overall mortality (5, 6). Chronic inflammation is implicated in sarcopenia as it interferes with the balance between muscle protein synthesis and breakdown (7).

Serum concentrations of IL-6, TNFα, and CRP have been shown to be elevated in sarcopenic elderly adults (14). A systematic review estimated that the prevalence of sarcopenia in RA is 31% (8). In RA, the presence of systemic, pro-inflammatory cytokines such as TNFα, IL-1, IL-6 and IFNγ promote skeletal muscle proteolysis and atrophy (9). Alterations in immune profile secondary to the autoimmune disease can influence skeletal muscle function (10). Furthermore, chronic inflammation, ageing, exercise and dietary habits also contribute to the development of sarcopenia in adults with inflammatory diseases such as rheumatoid arthritis (RA) (8, 9, 11, 12).

Juvenile idiopathic arthritis (JIA) is a chronic, autoimmune inflammatory disease (13) with poor outcomes in young adults even in the era of biologics (14). Current evidence suggests that similar to RA and ageing, sarcopenia and reduced bone mass are features of JIA in association with high disease activity (15-19). A recent study reported a significantly higher prevalence of severe sarcopenia of 33% in JIA when compared to 10% in ankylosing spondylitis and 18% in rheumatoid arthritis (20). Furthermore, specific subpopulations of lymphocytes such as memory B cells and effector memory T cells that re-express CD45RA (TEMRA), known for contributing to nonspecific inflammation, are notable at an increased frequency in chronic inflammatory conditions such as RA and ageing (21-23) and JIA (24).

However, our understanding of the relationship between skeletal muscle function, exercise and peripheral blood immune profile in young adults with JIA is not known.

Given the high prevalence of sarcopenia in the elderly, the SARC-F questionnaire, was developed and validated as a screening tool (25-28) and used pragmatically in people with musculoskeletal symptoms (29). We have recently described the potential utility of the SARC-F questionnaire in identifying people with JIA with sarcopenia and risk of falls (Manuscript in preparation). However, in JIA, the relationship between impaired muscle function, as assessed using SARC-F, and peripheral blood immune profile remains limited.

Furthermore, dietary and exercise habits are known to influence muscle function and/or immune profiles (11, 12, 30-37). Of direct relevance, a systematic review reported on the benefits of exercise on peripheral blood immune profile including B cells, T cells, NK cells and monocytes (36, 38-44) in addition to improving musculoskeletal function (45). Thus, exercise interventions and habits have the potential to restore peripheral blood immune profile and sarcopenia.

To this end, we explored the relationship between musculoskeletal function and immune profile, we compared the frequency of peripheral blood lymphocytes and monocytes in people with JIA categorised based on SARC-F score and exercise habits. We demonstrate that in people with JIA SARC-F scores relate to alterations in CD4 T cells and moderate exercise is associated with significantly lower frequency of TEMRA cells.

## Materials and Methods

### Participants

A total of 18 patients under the care of the Young Adult and Adolescent Rheumatology team at University College London Hospital participated in the study. All participants satisfied diagnostic criteria for JIA as defined by the International League of Associations for Rheumatology (46). Patients under the age of 18 years were excluded from the study. Data about current medications including Disease Modifying Anti Rheumatic Drugs, biologics and glucocorticoid use in the last six months was collected from electronic patient records.

### Musculoskeletal function questionnaire

Participants were invited to complete a short online questionnaire. The questionnaire comprised the SARC-F questionnaire and additional questions that included self-reported exercise habits. The SARC-F questionnaire (supplementary table 1) is a validated screening tool to identify the risk of sarcopenia (26). In this study, SARC-F scores were used as a surrogate measure of musculoskeletal function. Participants with a SARC-F score of four or more were considered to have musculoskeletal function impairment (SARC-F^hi^ +) or not impaired (SARC-F^hi^ -). Participants were also categorised according to the frequency of self-reported moderate exercise. Those performing moderate exercise more than once a week were categorised as the ‘exercise’ group (moderate exercise +) and the remaining were categorised as the ‘no exercise’ group (moderate exercise -).

### Flow cytometry

Peripheral blood samples were collected in VACUETTE tubes anticoagulated with lithium heparin and at least 10 million PBMCs were stored in liquid nitrogen before flow cytometry assessment using the BD LSRFortessa™ X-20 Cell Analyser. The flow cytometry data was analysed using the FlowJo™ version 10.10.0 Software.

### Statistical analysis

Statistical analysis and data visualisation was conducted using GraphPad Prism 10.2.3. Participant JIA10 was receiving treatment with belimumab, a monoclonal antibody targeting BAFF-R, contributing to low B cell counts. For this reason, JIA10 was excluded from all CD19+ B cell analysis. The Mann-Whitney U test was performed to test differences between lymphocyte subsets between the SARC-F^hi^+ and SARC-F^hi^-groups and the moderate exercise and no moderate exercise groups.

## Results

### Patient demographics and musculoskeletal function

Of the 18 participants, five were in the SARC-F^hi^+ group and 13 in the SARC-F^hi^-group. The median age of both the SARC-F^hi^+ and SARC-F^hi^-groups was 24 years. There were four females (80%) in the SARC-F^hi^+ group compared to five (38.46%) in the SARC-F^hi^-. SARC-F^hi^+ and SARC-F^hi^-subgroup demographics are detailed in Table 1.

**Table 1.** Demographic data of the SARC-F^hi^+ and SARC-F^hi^-subgroups.

| Demographics |  | SARC-F<sup>hi</sup>+ (n=5) | SARC-F<sup>hi</sup>- (n=13) |
| --- | --- | --- | --- |
| Age, years | Mean ± SD | 23±2.07 | 24.08±4.61 |
|  | Median | 24 | 24 |
| Sex (%) | Female | 4 (80.00%) | 5 (38.46%) |
| Height, cm | Mean ± SD | 167.75±8.54 | 170.17±13.53 |
|  | Median | 165 | 168 |
| Weight, kg | Mean ±SD | 69.72±5.90 | 69.92±14.32 |
|  | Median | 67 | 64.50 |
| JIA subtype (%) | Systemic arthritis | 0 | 2 (15.38%) |
|  | Oligoarthritis | 1 (20.00%) | 3 (23.08%) |
|  | Extended oligoarthritis | 0 | 2 (5.38%) |
|  | Polyarthritis | 3 (60.00%) | 2 (15.38%) |
|  | Psoriatic arthritis | 0 | 1 (7.69%) |
|  | Enthesis-related arthritis | 0 | 2 (15.38%) |
|  | Undifferentiated arthritis | 0 | 1 (7.69%) |
|  | Undifferentiated JIA with lupus | 1 (20.00%) | 0 |
| DMARD/Biologic therapy, % | Abatacept | 1 (20.00%) | 0 |
|  | Etanercept | 0 | 1 (7.69%) |
|  | Methotrexate | 1 (20.00%) | 1 (7.69%) |
|  | Infliximab | 2 (40.00%) | 4 (30.77%) |
|  | Tocilizumab | 1 (20.00%) | 4 (30.77%) |
|  | Adalimumab | 0 | 1 (7.69%) |
|  | Tofacinib | 0 | 1 (7.69%) |
|  | Belimumab | 0 | 1 (7.69%) |
| Oral or intravenous steroid use in last six months |  | 3 (60.00%) | 3 (23.08%) |

**Table 2.** Demographic data of the moderate exercise and no moderate exercise groups.

| Demographics |  | Moderate exercise (n=13) | No moderate exercise (n=5) |
| --- | --- | --- | --- |
| Age, years | Mean ± SD | 24±4.50 | 24±2.77 |
|  | Median | 23 | 25.00 |
| Sex (%) | Female | 6 (46.15%) | 3.00 (60.00%) |
| Height, cm | Mean ± SD | 170.67±12.66 | 167±10.23 |
|  | Median | 166.5 | 166.50 |
| Weight, kg | Mean ±SD | 72.28±13.03 | 63.58±8.49 |
|  | Median | 68.60 | 65.00 |

### Peripheral blood immune profile

Flow cytometry gating strategy is shown in Supplementary Figure 1. There were no significant differences in the frequency of CD19+CD3-B cells, CD19-CD3+ T cells and CD19-CD3-cells between the SARC-F^hi^+ and SARC-F^hi^-groups, shown in Figure 1 and Supplemental Table 4.

**Figure 1.**
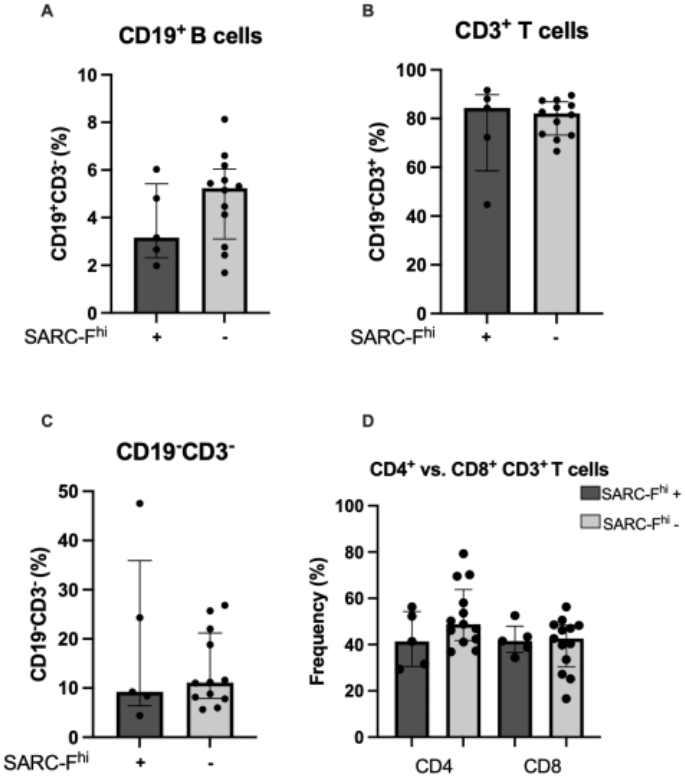
Peripheral blood B cells, T cells and NK cells based on SARC-F scores. Bar graphs represent the frequency of (A) CD19^+^CD3^−^B cells, (B) CD19^−^CD3^+^ T cells and (C) CD19^−^CD3^−^cells in the SARC-Fhi^+^and SARC-Fhi^−^groups. (D) shows the frequency of CD4^+^and CD8^+^cells within the CD19^−^CD3^+^ T-cell gate.

### B cells

JIA and exercise habits can influence the frequency of peripheral blood B cell subpopulations (47, 48). However, this study did not find any significant differences in B cell subpopulations (Figure 2 A - D).

**Figure 2.**
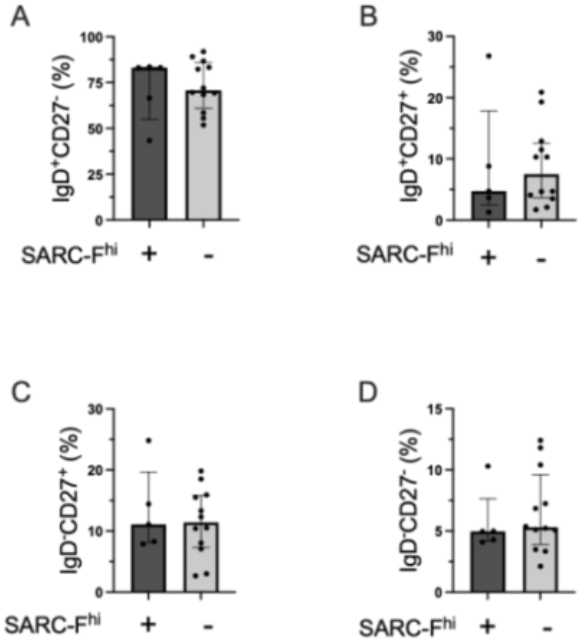
B cell subpopulations based on SARC-F scores. (A), IgD+CD27- B cells, (B) IgD+CD27+ B cells, (C) IgD-CD27+ B cells and D) IgD-CD27-B cells in SARC-F^hi^+ and SARC-F^hi^-groups.

### T cells

T cell profile changes with exercise habits and chronic inflammation in JIA (24, 40). Both the SARC-F^hi^+ and SARC-F^hi^-groups had similar frequencies of CD4+ and CD8+ T cells. The SARC-F^hi^+ group had 41.4% CD4+ and 41.5% CD8+ T cells. Similarly, the SARC-F^hi^-group have 48.80% CD4+ T cells and 42.6% CD8+ T cells (Figure 1D).

In the CD4+ population, the frequency of CD45RA-CD27+ central memory (CM) T cells was 45.6% in the SARC-F^hi^+ group and 45.5% in the SARC-F^hi^-group and the frequency of CD45A+CD27-TEMRA cells was low at 0.56% in the SARC-F^hi^+ and 0.19% in the SARC-F^hi^-groups. However, the SARC-F^hi^+ group had significantly lower CD45RA+CD27+ naïve T cells (26.6% v. 45.3%%, p = 0.0264) (Figure 2B) and higher CD45RA-CD27-effector memory (EM) T cells (11.90 v. 7.00%, p = 0.0350) (Figure 3D) (Supplementary table 4). There were no significant differences in the frequency of CD8+ T cell populations (figure 2E-H).

**Figure 3.**
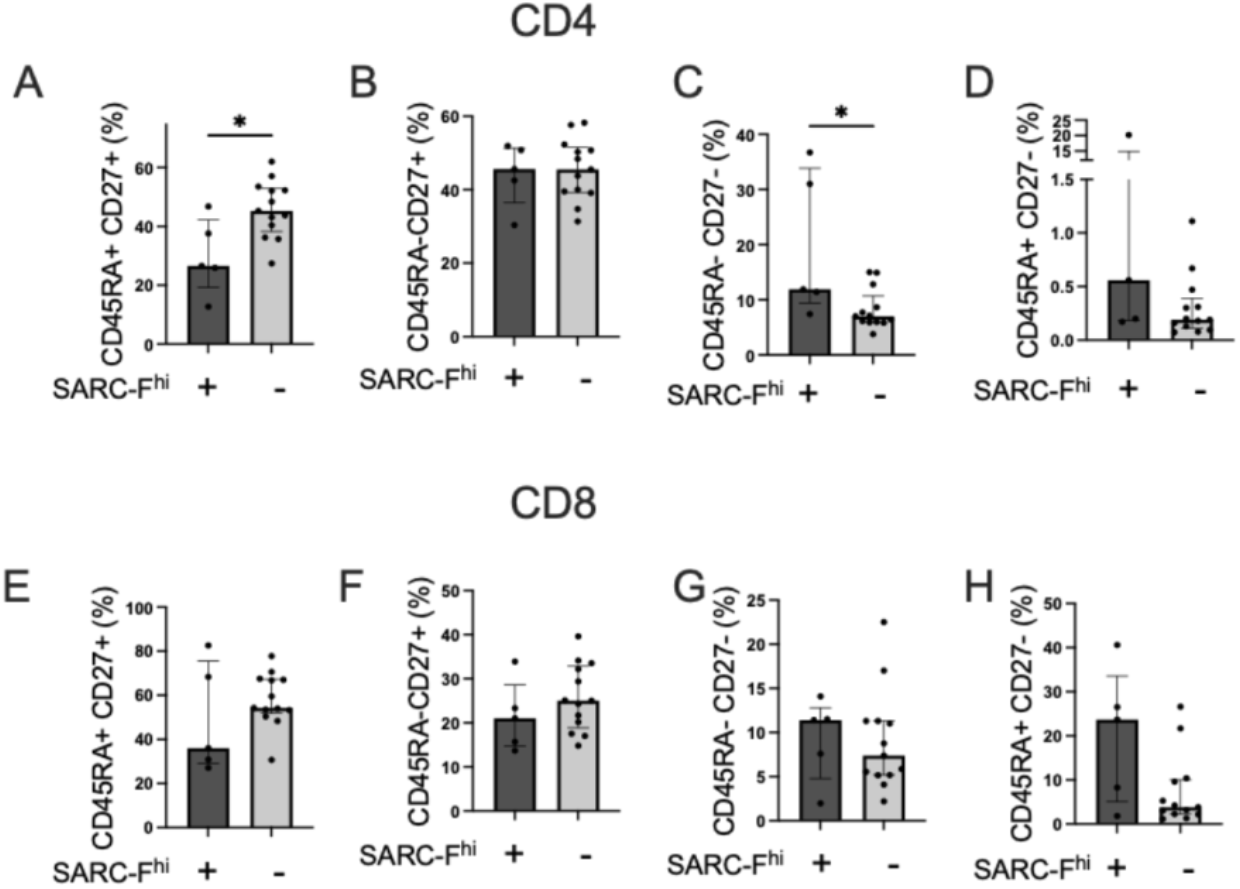
CD4 and CD8 T cell subpopulations in patients with JIA grouped based on SARC-F scores. Bar graphs represent the frequency of CD4^+^ T cells (A), naïve CD45RA^+^CD27^+^ T cells (B), central memory (CM) CD45RA^−^CD27^+^ T cells (C), effector memory (EM) CD45RA^−^CD27^−^T cells (D) and EMRA CD45RA^+^CD27^−^T cells in the SARC-Fhi^+^and SARC-Fhi^−^groups. * p < 0.05.

### NK cells

Chronic inflammation in JIA (49) and exercise habits influence NK cell frequency in peripheral blood (37). The frequency of CD56dimCD16+ NK cells was 31.5% in the SARC-F^hi^+ group and 57.2% in the SARC-F^hi^-group. There was a significant difference in the frequency of CD56-CD16+ NK cells with the SARC-F^hi^-group having a greater proportion (4.28%) as compared to the SARC-F^hi^+ group (2.6%) (p = 0.0194) (Figure 4A). The frequencies of the remaining CD56brightCD16+, CD56brightCD16-, CD56dimCD16+ and CD56dimCD16-groups were similar (Supplementary Table 5).

**Figure 4.**
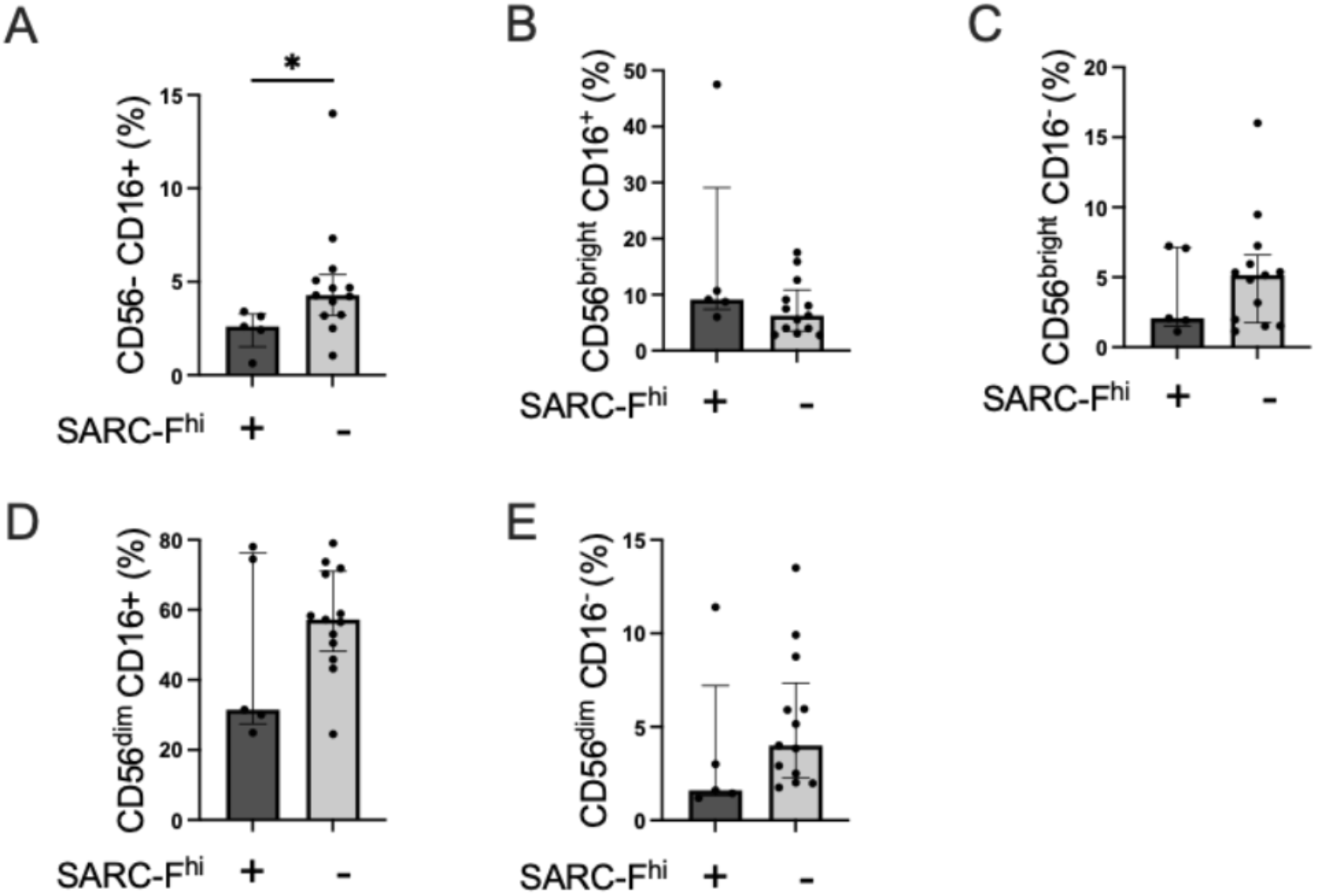
NK cell subpopulations based on SARC-F scores. Bar graphs represent the frequency of CD56^−^CD16^+^(A), CD56brightCD16^+^(B), CD56brightCD16^−^(C), CD56dimCD16^−^(D) and CD56dimCD16^+^(E) NK cells in the SARC-Fhi^+^and SARC-Fhi^−^groups. * p < 0.05.

### Moderate exercise

There were 13 participants in the exercise group and five in the no exercise group. The median ages in the exercise and no exercise groups were 23 years and 25 years, respectively. There were six females (46.15%) in the exercise group and three (60%) in the no exercise group (Table 4).

### Peripheral blood immune profile and exercise

There were similar frequencies of CD19+CD3-B cells, CD19-CD3+ T cells and CD19-CD3-cells (Supplementary Table 6).

### B cells

Alterations in B cell profile can occur in JIA and with exercise habits (47, 48). However, we did not find any significant differences in B cell subpopulations between the two groups categorised based on self-reported moderate exercise (Figure 5 A-D).

**Figure 5.**
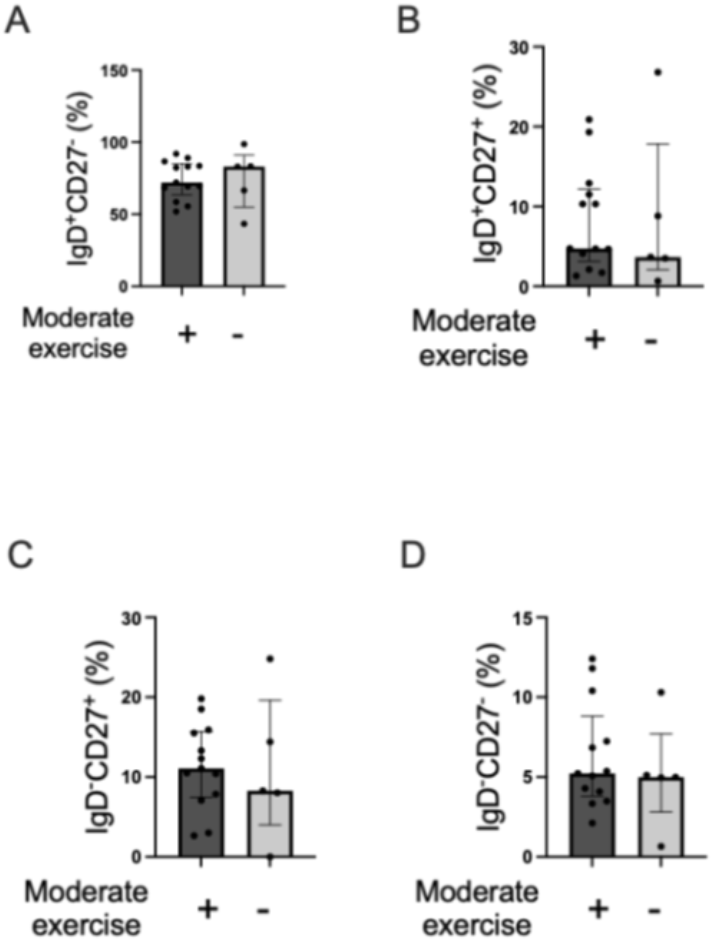
B cell subpopulations in patients with JIA grouped by self-reported exercise habits. Bar graphs represent the frequency of IgD^+^CD27^−^B cells (A), IgD^+^CD27^+^B cells (B), IgD^−^CD27^+^B cells (C) and IgD^−^CD27^−^B cells (D) in the Moderate + group (participants who perform moderate exercise) and Moderate - group (participants with no moderate exercise).

**Figure 6.**
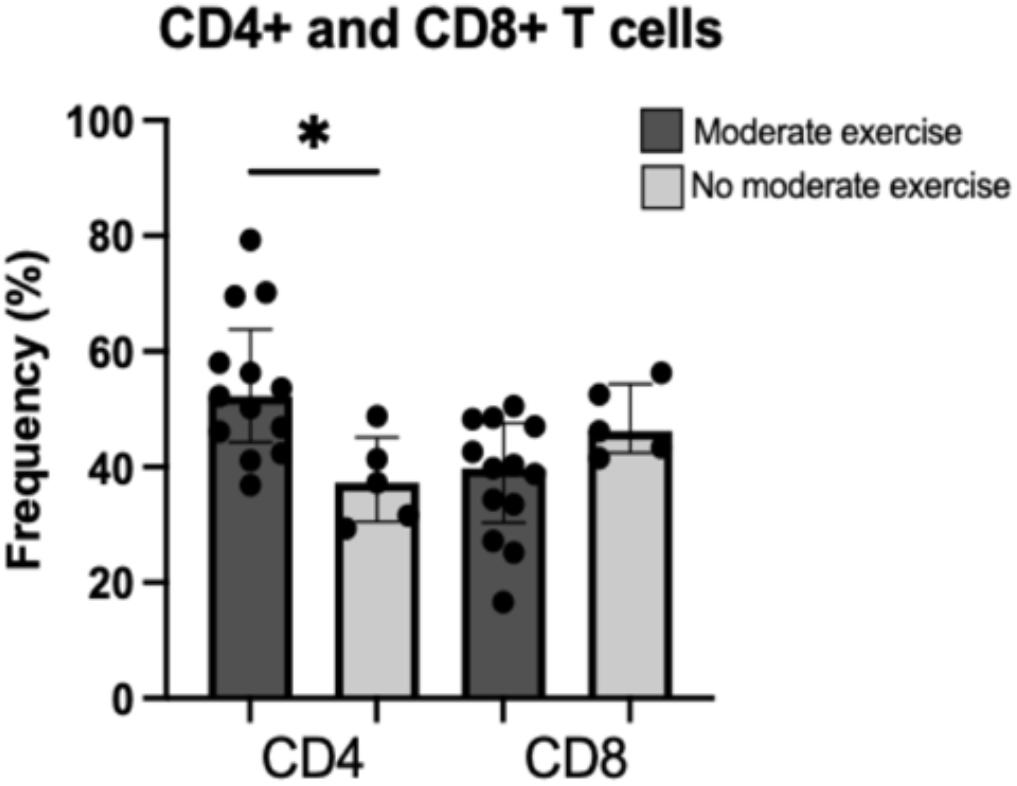
Frequency of CD4+ and CD8+ T cells affected by Moderate Exercise. Bar graphs represent the frequency of CD4^+^and CD8^+^ T cells in the Moderate^+^(exercise) and Moderate^−^(no exercise) groups. * p < 0.05.

### T cells

JIA and exercise habits can alter T cell profile (24, 40). The exercise group had significantly greater number of peripheral CD4+ T cells (52.30%) compared to the no exercise group (37.30%), (p = 0.0140), whereas no such differences between the groups were noted for CD8+ T cells, Supplementary Table 7.

There were no significant differences between the two groups in CD4+ T cell subpopulations (Figure 7 A – D). In the CD8+ T cell population, however, CD45RA+CD27-CD8+ TEMRA cells in the exercise group were three-fold lower compared to the no exercise group (10.40% vs. 3.17%, *p* = 0.0350). The frequencies of the remaining CD45RA+CD27+ naïve, CD45A-CD27+ CM, and CD45RA-CD27-EM CD8+ T cells were similar across groups (Figure 7 A-H, supplementary table 8). Therefore, the exercise group had a significantly lower frequency of senescence related CD8+ TEMRA cells.

**Figure 7.**
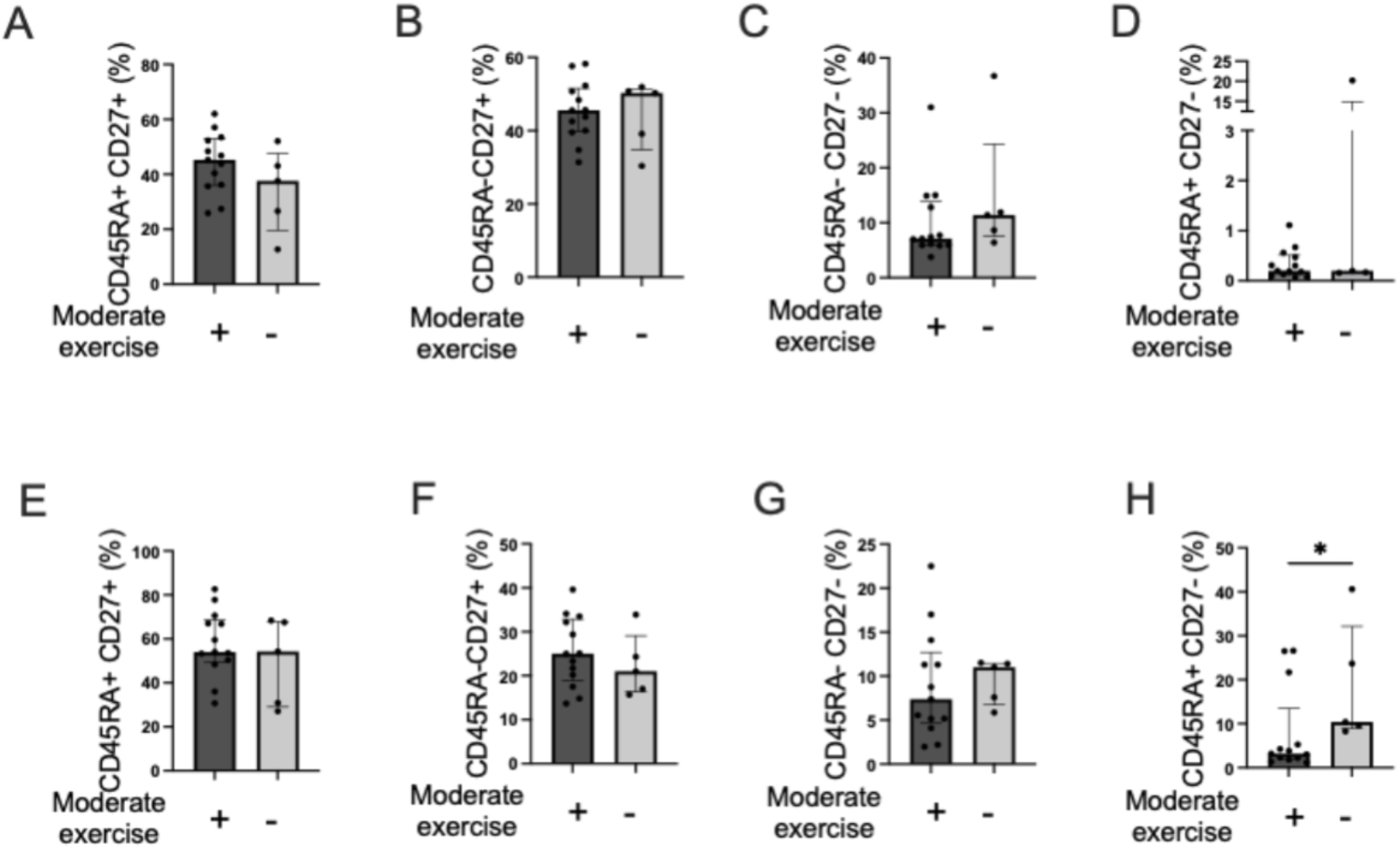
CD4 and CD8 T cell subpopulations in patients with JIA grouped based on self-reported exercise habits. Bar graphs represent the frequency of CD8^+^EMRA CD45RA^+^CD27^−^T cells (A), naïve CD45RA^+^CD27^+^ T cells (B), central memory (CM) CD45RA^−^CD27^+^ T cells (C), and effector memory (EM) CD45RA^−^CD27^−^T cells (D) in the Moderate^+^and Moderate^−^groups. * p < 0.05.

**Figure 8.**
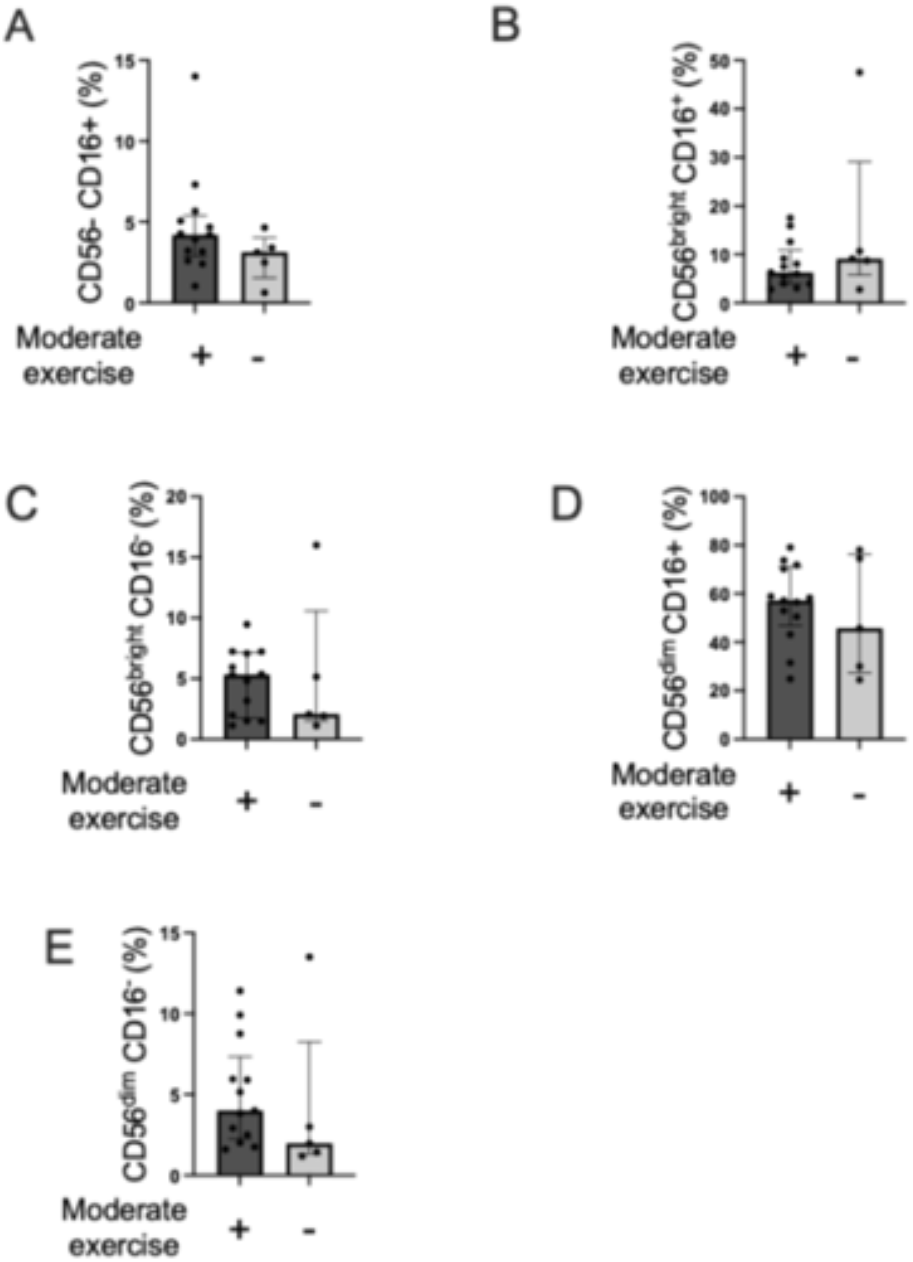
NK cell subpopulations in patients with JIA grouped based on self-reported exercise habits. Bar graphs represent the frequency of CD56^−^CD16^−^ NK cells (A), CD56brightCD16^+^ NK cells (B), CD56brightCD16^−^ NK cells (C), CD56dimCD16^+^ NK cells (D) and CD56dimCD16^−^ NK cells (E) in the Moderate + group (participants who perform moderate exercise) and Moderate - group (participants with no moderate exercise).

### NK cells

Previous reports significant alterations in NK cells in both JIA and exercise (37, 50). However, we found no significant differences in NK cell subpopulations based on self-reported exercise.

## Discussion

Considering the dynamic interactions between the skeletal muscle and the immune system in health and disease (1, 10, 51), a key aim of this study was to initiate the exploration of the relationship between musculoskeletal function and peripheral blood immune profile in young people with JIA.

## Limitations

Our study has several limitations. Fist, this was a single centre observational study that was not powered to account for the potential impact of factors other than the diagnosis of JIA and exercise habits such as medications including corticosteroids and biologics that could influence peripheral blood immune profile. Second, participants were recruited in outpatient settings and did not have high disease activity, therefore, these findings may not apply to patients with acute high disease activity. Third, we have not evaluated immune profile of internal organs such as the joint and muscle. Fourth, SARC-F score, but not comprehensive measurements of muscle mass, strength etc., was used as a measure of musculoskeletal function. Finally, the cryopreservation process of PBMCs may differentially affect cell viability and enrich for certain cell types. The discussion that follows considers these limitations.

Alterations in immune profile during ageing (52) and exercise are well described (40). In this study, the SARC-F^hi^+ group had fewer peripheral naïve CD4+ T cells as compared to the SARC-F^hi^-group and trends towards an association between musculoskeletal function and increased frequency of CD4+ and CD8+ TEMRA cells were notable. TEMRA cells are terminally differentiated effector memory T cells with the capacity to secrete cytotoxic and inflammatory molecules and cause nonspecific tissue damage accumulate in ageing (52) and JIA (24) and exercise is associated with reduced frequency (38, 40). These findings support previous reports of T cell immunosenescence in JIA (24) with add evidence of a relationship with poor musculoskeletal function, thereby providing the rationale for modifying lifestyle such as exercise (40).

Our study is the first to report, in JIA on the relationship between peripheral CD4+ T cells and musculoskeletal functional impairment. Participants with musculoskeletal functional impairment had fewer circulating CD56-CD16+ NK cells compared to those without. CD56-CD16+ NK cells have primarily been studied in the context of chronic infectious diseases (53). There is a lack of literature on the role of CD56-CD16+ NK cells in JIA in the context of interactions with skeletal muscle. This study raises the question of how the bidirectional relationship between skeletal muscle and NK cells may change with skeletal muscle functionality in chronic inflammation.

This study has suggested that exercise can impact peripheral T cells in JIA. The exercise group had significantly increased total CD4+ T cells, however, CD4+ T cell subpopulations did not differ between the groups. The difference in total CD8+ T cell proportion was insignificant, however, the exercise group had three-fold lower CD8+ TEMRA cells compared to the non-exercise group. Few studies have examined the impact of exercise on the peripheral immune profile in JIA patients, especially those with poor musculoskeletal function. Our findings indicate that participants with poor musculoskeletal function may exhibit immune profile features similar to ageing adults, particularly in naïve and TEMRA cell frequencies (54). Collectively, these observations suggest that exercise may influence the peripheral T cell pool in a manner that slows or prevents TEMRA cell accumulation in JIA..

## Future perspectives

This study highlights significant variations in immune profiles related to musculoskeletal function and exercise in young adults with JIA. Future studies overcoming the limitations of our study are warranted to improve understanding of sarcopenia and peripheral blood immune profile in JIA, particularly focusing on moderate exercise’s impact on total CD4^+^populations and CD8^+^ TEMRA cells. Furthermore, investigating other factors such as dietary habits and vitamin D status may inform strategies to promote skeletal muscle function and restore a favourable immune profile.

## Conclusions

Sarcopenia is a disorder of skeletal muscle affecting not only older adults but also young adults with inflammatory arthritis. This study demonstrates that JIA patients with musculoskeletal functional impairment have fewer naïve and greater effector memory CD4^+^ T cells, suggesting a relationship between musculoskeletal dysfunction and immunosenescence in this cohort. Further research into immune–muscle interactions in JIA may advance efforts to reverse sarcopenia and restore immune homeostasis.

## Acknowledgements

VR received funding support for this work from MRC-CARP Fellowship (MR/T024968/1), Versus Arthritis and UCL Division of Medicine Fellowship (2024). DS and SS received funding support for this work from the MRC CAPE grant.

