## Supplementary figures and images for "Relationship between SARC-F score and Peripheral Blood Lymphocytes in Juvenile Idiopathic Arthritis: An Observational Pilot Study"

### Supplemental Figure 1

#### Supplementary Material

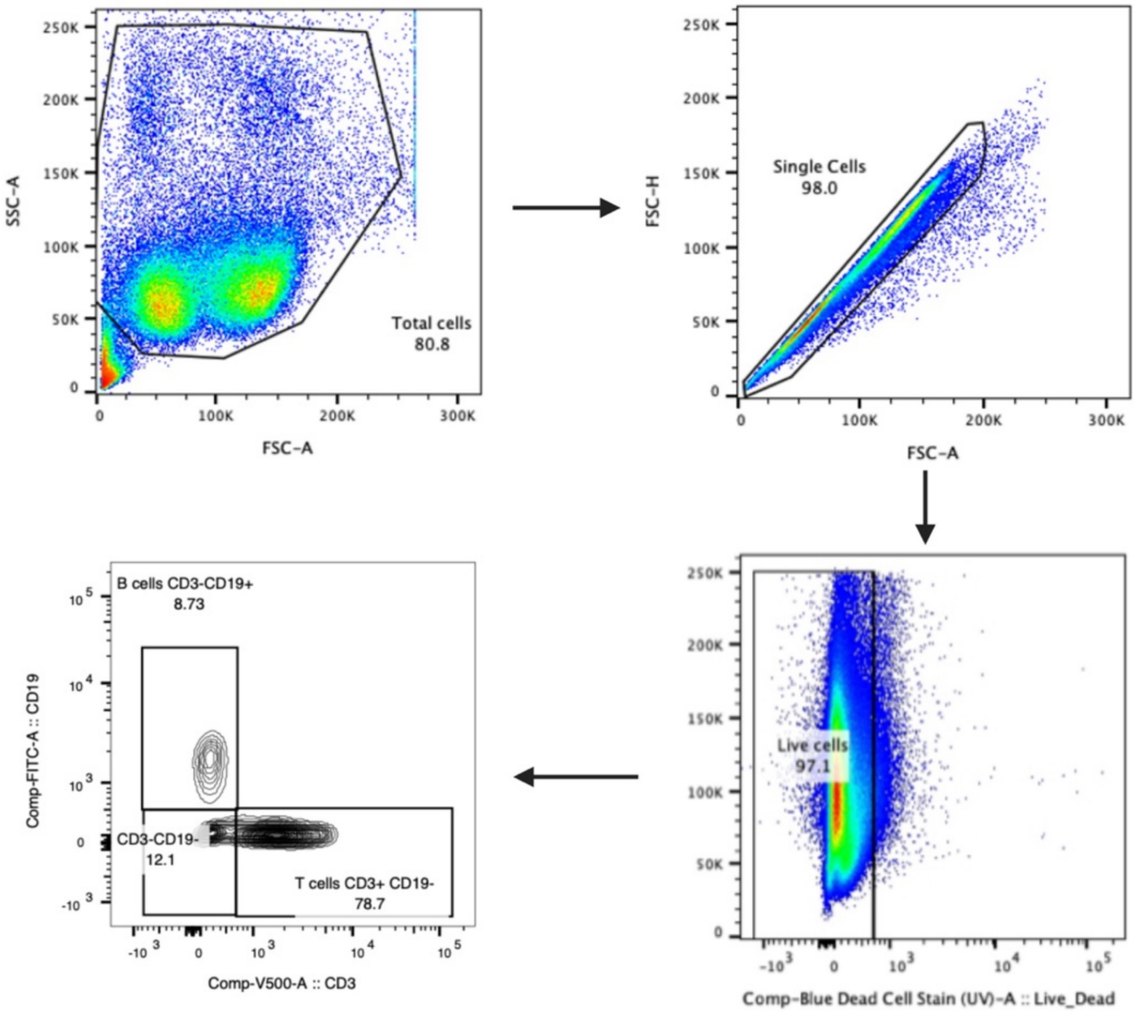


**Supplementary Figure 1.** *Gating Strategy performed on stained PBMCs.*
