## Supplemental Figure 2 for "Relationship between SARC-F score and Peripheral Blood Lymphocytes in Juvenile Idiopathic Arthritis: An Observational Pilot Study"

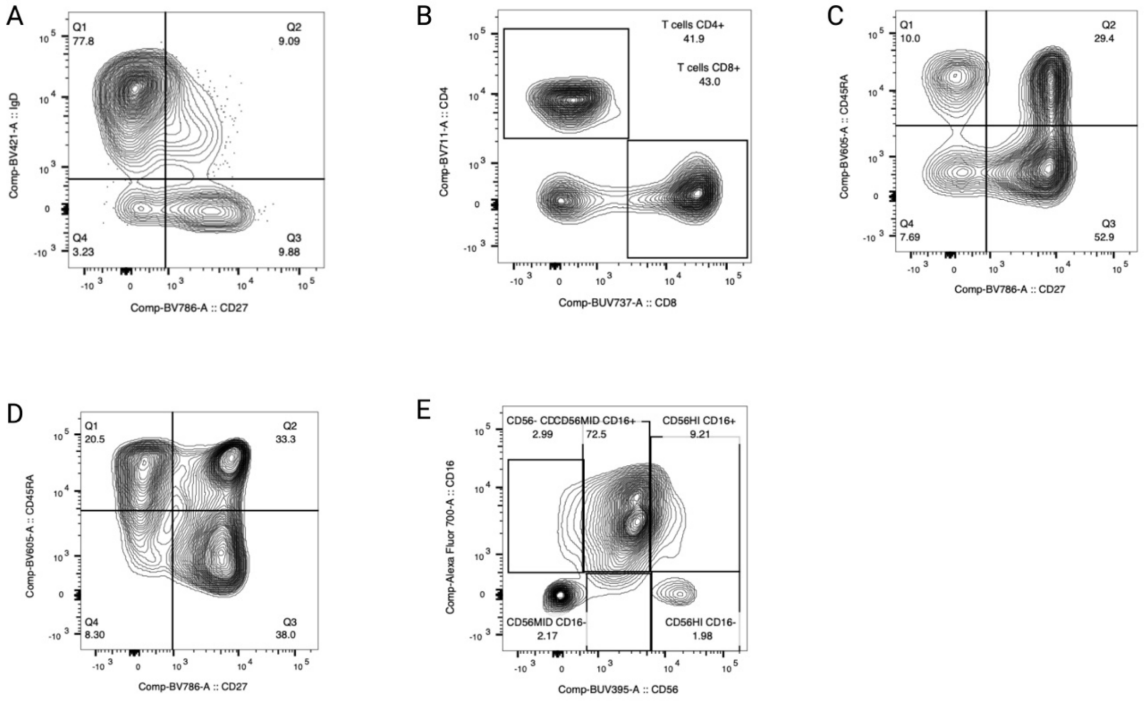


**Supplementary Figure 2.** *Lymphocyte subset gating.* (A) B cell subsets performed on the CD19+CD3- gate. (B) CD4+ and CD8+ T cell populations, performed in the CD19-CD3+ gate. (C) the gating performed on the CD4+ T cell gate. (D) represents the gating performed on the CD8+ T cell gate. (E) the gating performed on the CD19-CD3- cells.
