## Supplemental Figure 3 for "Relationship between SARC-F score and Peripheral Blood Lymphocytes in Juvenile Idiopathic Arthritis: An Observational Pilot Study"

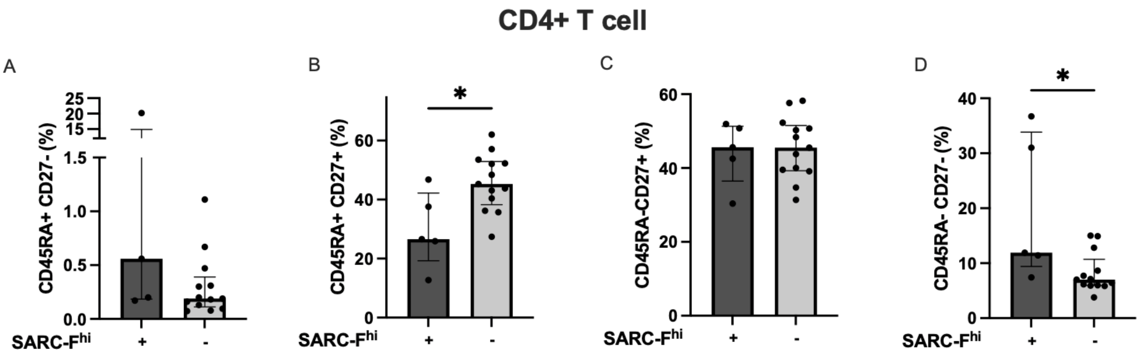


**Supplementary Figure 3.** *Frequency of CD4+ T cell subsets in the SARC-$F^{hi}$+ and SARC-$F^{hi}$- groups*. CD4$^{+}$ T-cell subpopulations are shown in bar graphs: (A) CD45RA$^{+}$CD27$^{-}$; (B) CD45RA$^{+}$CD27$^{+}$; (C) CD45RA$^{-}$CD27$^{+}$; and (D) CD45RA$^{-}$CD27$^{-}$. These subsets are compared between the SARC-Fhi$^{+}$ and SARC-Fhi$^{-}$ groups.
