## Supplemental Table 1 for "Relationship between SARC-F score and Peripheral Blood Lymphocytes in Juvenile Idiopathic Arthritis: An Observational Pilot Study"

**Supplementary Table 1.** *SARC-F questionnaire* (10)

| **Question** | **Response** | **Score** |
|---|---|---|
| How much difficulty to you have in lifting 10 pounds? | None | 0 |
| | Some | 1 |
| | A lot or unable | 2 |
| How much difficulty do you have walking across a room? | None | 0 |
| | Some | 1 |
| | A lot, use aids, or unable | 2 |
| How much difficulty do you have transferring from a chair or bed? | None | 0 |
| | Some | 1 |
| | A lot or unable without help | 2 |
| How many times have you fallen in the past year? | None | 0 |
| | Less than three | 1 |
| | Four or more | 2 |
