## Supplemental Table 2 for "Relationship between SARC-F score and Peripheral Blood Lymphocytes in Juvenile Idiopathic Arthritis: An Observational Pilot Study"

**Supplementary Table 2.** *Peripheral lymphocyte phenotypes*

| Cell type | Phenotype |
|---|---|
| CD19+ B cells | IgD+CD27- naïve |
| | IgD+CD27+ unswitched memory |
| | IgD-CD27+ switched memory |
| | IgD-CD27- double negative (DN) |
| CD3+ CD4+ T cells<br>CD3+ CD8+ T cells | CD45RA+CD27+ naïve |
| | CD45RA-CD27+ central memory (CM) |
| | CD45RA-CD27- effector memory (EM) |
| | CD45RA+CD27- TEMRA |
| CD19-CD3- NK cells | CD56-CD16+ |
| | CD56dimCD16+ |
| | CD56brightCD16+ |
| | CD56dimCD16- |
| | CD56brightCD16- |
