## Supplemental Table 3 for "Relationship between SARC-F score and Peripheral Blood Lymphocytes in Juvenile Idiopathic Arthritis: An Observational Pilot Study"

**Supplementary Table 3.** Antibodies and fluorophores used for identification of peripheral lymphocytes

| **Antigen** | **Fluorophore** | **Manufacturer** | **Clone** | **Lot** | **Volume (µl)** |
|---|---|---|---|---|---|
| Live-Dead | Zombie UV | BioLegend | N/A | B369909 | 0.5 |
| CD19 | FITC | BioLegend | HIB19 | B274550 | 1 |
| IgD | Brilliant Violet 421 | BioLegend | IA6-2 | B384772 | 1 |
| CD27 | Brilliant Violet 786 | BD Biosciences | L128 | 3044374 | 1 |
| CD3 | V500 | BD Biosciences | UCHT1 | 0107958 | 1 |
| CD4 | Brilliant Violet 711 | BioLegend | RPA-T4 | B329655 | 1 |
| CD8 | BUV737 | BD Biosciences | SK1 | 3212826 | 1 |
| CD45RA | Brilliant Violet 605 | BioLegend | HI100 | B281355 | 1 |
| CD56 | BUV395 | BD Biosciences | NCAM16.2 | 3132991 | 1 |
| CD16 | Alexa Flour 700 | BioLegend | 3G8 | B266048 | 1 |
