## Supplemental Table 4 for "Relationship between SARC-F score and Peripheral Blood Lymphocytes in Juvenile Idiopathic Arthritis: An Observational Pilot Study"

**Supplementary Table 4.** *Frequency of CD19+CD3- B cells, CD19-CD3+ T cells and CD19-CD3- cells in the SARC-F$^{hi}$+ and SARC-F$^{hi}$- groups.* Analysis was performed after gating for total cell population, single cells and live cells. P values were calculated using the Mann-Whitney test and significance was denoted at p < 0.05. Participant JIA10, highlighted with an asterisk, was excluded from analysis due to systemic treatment with BAFF inhibitor belimumab.

| | **Participant study number** | **Frequency (%)** | | |
|---|---|---|---|---|
| | | CD19+ CD3- | CD19-CD3+ | CD19-CD3- |
| **SARC-F$^{hi}$ +** | JIAPS07 | 1.98 | 88 | 9.23 |
| | JIAPS35 | 3.16 | 91.6 | 4.38 |
| | JIA01 | 2.66 | 72.3 | 24.3 |
| | JIA12 | 4.81 | 84.4 | 8.38 |
| | JIAPS28 | 6.03 | 44.7 | 47.5 |
| | **Median** | 3.160 | 84.40 | 9.23 |
| | **Mean** | 3.73 | 76.2 | 18.76 |
| | **Standard deviation** | 1.66 | 19.05 | 17.76 |
| **SARC-F$^{hi}$ -** | JIAPS18 | 6.19 | 84.5 | 7.8 |
| | JIAPS21 | 5.15 | 87.6 | 5.64 |
| | JIAPS30 | 4.47 | 82.7 | 11.6 |
| | JIA02 | 5.44 | 87.4 | 5.97 |
| | JIA03 | 5.32 | 66.6 | 26.8 |
| | JIA04 | 8.13 | 78.7 | 11.2 |
| | JIA05 | 5.57 | 73.6 | 18.8 |
| | JIA06 | 6.6 | 81.5 | 8.83 |
| | JIA07 | 2.76 | 85.4 | 11 |
| | JIA08 | 4.13 | 73.1 | 22 |
| | JIA09 | 2.43 | 71.2 | 25.7 |
| | JIA10* | 0.067 | 96.2 | 3.21 |
| | JIA11 | 1.69 | 89.5 | 8.13 |
| | **Median** | 5.235 | 82.10 | 11.10 |
| | **Mean** | 4.82 | 80.15 | 13.62 |
| | **Standard deviation** | 1.85 | 7.44 | 7.644 |
| **P value (<0.05)** | | 0.3284 | 0.8788 | 0.9593 |
