## Supplemental Table 5 for "Relationship between SARC-F score and Peripheral Blood Lymphocytes in Juvenile Idiopathic Arthritis: An Observational Pilot Study"

**Supplementary Table 5.** *Frequency of CD4+ T cell subsets in the SARC-F$^{hi}$+ and SARC-F$^{hi}$- groups.* Analysis was performed after gating on total cell population, single cells, live cells, $CD19^{-}CD3^{+}$ cells and $CD4^{+}CD8^{-}$ cells. P values were calculated using the Mann–Whitney test, and asterisks denote $p < 0.05$.

| | Participant study number | Frequency of CD4+ T cell subsets (%) | | | |
|---|---|---|---|---|---|
| | | CD45RA+ CD27- | CD45RA+ CD27+ | CD45RA- CD27+ | CD45RA- CD27- |
| **SARC-F$^{hi}$ +** | JIAPS07 | 9.56 | 26.6 | 51.9 | 11.9 |
| | JIAPS35 | 0.2 | 46.8 | 45.6 | 7.4 |
| | JIA01 | 20.2 | 12.7 | 30.4 | 36.7 |
| | JIA12 | 0.56 | 25.9 | 42.5 | 31 |
| | JIAPS28 | 0.17 | 37.6 | 50.8 | 11.4 |
| | **Median** | 0.5600 | 26.60 | 45.60 | 11.90 |
| | **Mean** | 6.14 | 29.92 | 44.24 | 19.68 |
| | **Standard deviation** | 8.82 | 12.92 | 8.63 | 13.21 |
| **SARC-F$^{hi}$ -** | JIAPS18 | 0.47 | 62 | 31.4 | 6.15 |
| | JIAPS21 | 0.16 | 52.1 | 39.1 | 8.65 |
| | JIAPS30 | 0.078 | 27.4 | 57.6 | 14.9 |
| | JIA02 | 1.11 | 57.1 | 34.7 | 7.11 |
| | JIA03 | 0.67 | 53.5 | 40 | 5.86 |
| | JIA04 | 0.3 | 36.2 | 50.7 | 12.8 |
| | JIA05 | 0.18 | 45.3 | 39.5 | 15 |
| | JIA06 | 0.13 | 35.7 | 58.2 | 5.99 |
| | JIA07 | 0.31 | 40.4 | 52.3 | 7 |
| | JIA08 | 0.19 | 48.4 | 45.5 | 5.9 |
| | JIA09 | 0.095 | 43.8 | 48.4 | 7.72 |
| | JIA10 | 0.2 | 43.1 | 50.3 | 6.41 |
| | JIA11 | 0.072 | 52.3 | 43.8 | 3.79 |
| | **Median** | 0.1900 | 45.30 | 45.50 | 7.000 |
| | **Mean** | 0.31 | 45.95 | 45.50 | 8.25 |
| | **Standard deviation** | 0.30 | 9.62 | 8.36 | 3.63 |
| **P value (<0.05)** | | 0.0983 | 0.0264* | >0.9999 | 0.0350* |
