## Supplemental Table 6 for "Relationship between SARC-F score and Peripheral Blood Lymphocytes in Juvenile Idiopathic Arthritis: An Observational Pilot Study"

**Supplementary Table 6.** *Frequency of NK cell subsets in the SARC-F$^{hi}$+ and SARC-F$^{hi}$- groups.* Analysis was performed after gating on total cell population, single cells, live cells and CD19$^{-}$CD3$^{-}$ cells. P values were calculated using the Mann–Whitney test, and significance was denoted at $p < 0.05$. Asterisks denote statistically significant p values.

| | **Participant study number** | **Frequency of NK cell subsets (%)** | | | | |
|---|---|---|---|---|---|---|
| | | CD56- CD16+ | CD56bright CD16+ | CD56bright CD16- | CD56dim CD16+ | CD56dim CD16- |
| **SARC-F$^{hi}$ +** | JIAPS07 | 3.4 | 8.73 | 2.08 | 74.5 | 3.01 |
| | JIAPS35 | 2.42 | 9.16 | 7.22 | 31.5 | 1.61 |
| | JIA01 | 3.14 | 10.7 | 1.12 | 78 | 1.19 |
| | JIA12 | 2.6 | 6.03 | 7.07 | 24.9 | 11.4 |
| | JIAPS28 | 0.63 | 47.5 | 1.92 | 30 | 1.44 |
| | **Median** | 2.600 | 9.160 | 2.080 | 31.50 | 1.610 |
| | **Mean** | 2.438 | 16.42 | 3.882 | 47.78 | 3.730 |
| | **Standard deviation** | 1.086 | 17.45 | 3.001 | 26.13 | 4.346 |
| **SARC-F$^{hi}$ -** | JIAPS18 | 4.22 | 3.06 | 5.94 | 53 | 5.9 |
| | JIAPS21 | 2.5 | 9.09 | 16 | 24.5 | 1.98 |
| | JIAPS30 | 3.23 | 15.9 | 5.38 | 57.2 | 5.95 |
| | JIA02 | 5.68 | 5.52 | 7.24 | 56.5 | 5.16 |
| | JIA03 | 5.07 | 2.83 | 1.14 | 73.7 | 4 |
| | JIA04 | 4.28 | 4.02 | 4.83 | 50.5 | 9.92 |
| | JIA05 | 14 | 12.6 | 1.97 | 58.3 | 2.5 |
| | JIA06 | 7.32 | 17.5 | 5.34 | 43.2 | 3.84 |
| | JIA07 | 3.18 | 6.25 | 9.48 | 58.9 | 8.75 |
| | JIA08 | 4.68 | 3.97 | 1.53 | 70.2 | 2.92 |
| | JIA09 | 1.04 | 8 | 1.5 | 79 | 1.75 |
| | JIA10 | 4.65 | 2.78 | 5.15 | 45.8 | 13.5 |
| | JIA11 | 3.94 | 7.49 | 3.18 | 71.9 | 2.02 |
| | **Median** | 4.280 | 6.250 | 5.150 | 57.20 | 4.000 |
| | **Mean** | 4.07 | 7.616 | 5.283 | 57.13 | 5.245 |
| | **Standard Deviation** | 3.130 | 4.942 | 4.061 | 14.71 | 3.570 |
| **P value (<0.05)** | | 0.0194* | 0.1433 | 0.6331 | 0.6631 | 0.1433 |
