## Supplemental Table 8 for "Relationship between SARC-F score and Peripheral Blood Lymphocytes in Juvenile Idiopathic Arthritis: An Observational Pilot Study"

**Supplementary Table 8**. *Frequency of CD4+ and CD8+ T cells in the moderate exercise and no moderate exercise groups.* Analysis was conducted after gating on total cell population, single cells, live cells and $CD19^{-}CD3^{+}$ T cells. P values were calculated using the Mann–Whitney test, and significance was denoted at $p < 0.05$. Asterisks denote significant p values.

| | **Participant study number** | **Frequency of T cell subtypes** | |
|---|---|---|---|
| | | CD4 | CD8 |
| **Moderate exercise** | JIAPS35 | 56.3 | 38.8 |
| | JIA12 | 52.3 | 34.4 |
| | JIAPS18 | 46.1 | 48.5 |
| | JIAPS30 | 41.1 | 50.5 |
| | JIA02 | 46.9 | 47 |
| | JIA03 | 70.2 | 25.2 |
| | JIA04 | 36.9 | 48.3 |
| | JIA05 | 42.4 | 39.8 |
| | JIA06 | 58 | 33.5 |
| | JIA07 | 50.2 | 42.6 |
| | JIA08 | 69.5 | 27.2 |
| | JIA09 | 53.6 | 40.4 |
| | JIA11 | 79.3 | 16.6 |
| | Mean | 54.06 | 37.91 |
| | Median | 52.30 | 39.80 |
| | Standard deviation | 12.54 | 10.23 |
| **No moderate exercise** | JIAPS07 | 41.4 | 43.4 |
| | JIAPS21 | 48.8 | 46.2 |
| | JIAPS28 | 31.6 | 41.5 |
| | JIA01 | 29.4 | 52.5 |
| | JIA10 | 37.3 | 56.3 |
| | Mean | 37.70 | 47.98 |
| | Median | 37.30 | 46.20 |
| | Standard deviation | 7.797 | 6.241 |
| **P value** | | 0.0140* | 0.0593 |
