## Supplemental Table 9 for "Relationship between SARC-F score and Peripheral Blood Lymphocytes in Juvenile Idiopathic Arthritis: An Observational Pilot Study"

**Supplementary Table 9.** *Frequency of CD8+ T cell subsets in the moderate and no moderate exercise groups.* Analysis was performed after gating on total cell population, single cells, live cells, $CD19^{-}CD3^{+}$ cells and $CD8^{+}CD4^{-}$ cells. P values were calculated using the Mann–Whitney test, and significance was denoted at $p < 0.05$. Asterisks denote significant p values.

| | **Participant study number** | **CD8 Frequency (%)** | | | |
|---|---|---|---|---|---|
| | | CD45RA+ CD27- | CD45RA+ CD27+ | CD45RA- CD27+ | CD45RA- CD27- |
| **Moderate Exercise** | JIAPS35 | 1.8 | 82.6 | 13.7 | 1.96 |
| | JIA12 | 26.5 | 36 | 23.3 | 14.1 |
| | JIAPS18 | 2.28 | 77.8 | 14.8 | 5.16 |
| | JIAPS30 | 1.15 | 48.4 | 33.5 | 17 |
| | JIA02 | 26.6 | 30.7 | 20.2 | 22.5 |
| | JIA03 | 5.26 | 54 | 29.4 | 11.3 |
| | JIA04 | 3.8 | 67 | 25.1 | 4.09 |
| | JIA05 | 3.02 | 66.8 | 25 | 5.18 |
| | JIA06 | 4.23 | 59.5 | 34.1 | 2.19 |
| | JIA07 | 21.7 | 53.5 | 17.5 | 7.39 |
| | JIA08 | 3.17 | 53.3 | 32.2 | 11.3 |
| | JIA09 | 1.32 | 50.4 | 39.6 | 8.76 |
| | JIA11 | 2.17 | 70.6 | 21.7 | 5.55 |
| | Mean | 7.923 | 57.74 | 25.39 | 8.960 |
| | Median | 3.170 | 54.00 | 25.00 | 7.390 |
| | Standard deviation | 9.832 | 15.15 | 7.970 | 6.102 |
| **No moderate exercise** | JIAPS07 | 23.7 | 30.9 | 33.9 | 11.5 |
| | JIAPS21 | 10.4 | 54.3 | 24.3 | 11 |
| | JIAPS28 | 8.29 | 68.4 | 15.7 | 7.6 |
| | JIA01 | 40.6 | 27.1 | 21 | 11.4 |
| | JIA10 | 9.6 | 67.5 | 17 | 5.86 |
| | Mean | 18.52 | 49.64 | 22.38 | 9.472 |
| | Median | 10.40 | 54.30 | 21.00 | 11.00 |
| | Standard deviation | 13.82 | 19.70 | 7.278 | 2.584 |
| **P value** | | 0.0350* | 0.7028 | 0.5028 | 0.4307 |
